# Integrative multiomics profiling of cortical brain organoids reveals a druggable alternative splicing program in schizophrenia

**DOI:** 10.64898/2026.07.06.736738

**Authors:** Ibrahim A. Akkouh, Jordi Requena Osete, Attila Szabo, Vidar M. Steen, Anja Torsvik, Espen Molden, Nadine Parker, Elise Koch, Michael Ziller, Ole A. Andreassen, Kevin S. O’Connell, Srdjan Djurovic

## Abstract

Clozapine (CLZ) is the only available pharmacological option for treatment resistant schizophrenia (TRS), but its use is limited due to adverse drug reactions and potential cytotoxicity. Despite decades of research, the precise mechanisms of action of CLZ in the human brain remain poorly understood. To address this, we derived cortical brain organoids from a large cohort of schizophrenia (SCZ) patients and healthy controls and employed a comprehensive multiomics strategy to dissect the cellular mechanisms of long-term CLZ exposure of up to 24 weeks. We uncovered a SCZ-specific and metabolism-independent alternative splicing program that was amenable to non-toxic CLZ treatment. In-depth analysis revealed a key role of exon skipping and intron retention in glutamatergic neurons. This program was further found to recapitulate disease mechanisms in primary human brain tissue and capture splicing-mediated genetic signals of SCZ risk. These findings highlight alternative splicing as a promising avenue for therapeutic developments in TRS.

## MAIN

Around a quarter of patients with schizophrenia (SCZ) do not respond adequately to conventional antipsychotic medication^1^. This clinical picture is known as treatment-resistant SCZ (TRS) and is associated with increased symptom severity, poorer neurocognitive functioning, and reduced quality of life^2^. TRS risk is influenced by common genetic variation^3, 4^, and although multiple neurobiological perturbations have been proposed^5^, the mechanistic underpinnings are poorly understood. As a consequence, available treatment options for TRS are severely restricted.

The second-generation antipsychotic clozapine (CLZ) is currently the only evidence-based and approved pharmacological option for TRS^6^. However, despite its superior efficacy, CLZ is underutilized due to adverse drug reactions and potential cytotoxicity, necessitating strict hematological monitoring^7^. In addition, CLZ has a prolonged adequate trial period, requiring at least 12 weeks of treatment to achieve 40-50% response rate^8^, which further undermines the full potential of the drug. The receptor binding profile of CLZ is highly complex, with unique actions at muscarinic and glutamatergic receptors^9^. While these characteristics have been suggested to mediate the clinical superiority of CLZ, its exact mechanisms of action in the human brain, especially intracellularly downstream of the receptor level, remain obscure^9, 10^. Better insight into the molecular mechanisms of CLZ treatment may help to discover novel therapeutical targets and replenish the repertoire of available treatment options for TRS.

Induced pluripotent stem cell (iPSC) systems enable high-fidelity modeling of CLZ mechanisms in the human brain. Previous studies have either investigated two-dimensional neural cultures or included a relatively small number of iPSC lines, and CLZ administration has not been extended beyond two weeks of exposure^11–14^. Moreover, CLZ undergoes a genetically influenced metabolism by cytochrome P450 (CYP) enzymes within the liver as well as the brain^15, 16^, but this important aspect has not been investigated.

Here we address these issues by deriving three-dimensional human cortical brain organoids (hCOs) from a large (n = 28) and genetically stratified cohort of iPSC lines from SCZ patients and healthy control (CTRL) donors. We employ an integrated multiomics analysis strategy to comprehensively assess the molecular mechanisms of action of CLZ after long-term exposure for up to 24 weeks. We identify a cell type-specific and druggable alternative splicing program in SCZ which is independent of CLZ metabolism. We show that this program recapitulates disease mechanisms in primary human brain tissue and captures the splicing-mediated genetic signal of SCZ risk.

## RESULTS

### Concordance of transcriptomic and proteomic hCO profiles

We derived hCOs from a well-characterized and previously described^17, 18^ cohort of genetically stratified SCZ (n = 14) and CTRL (n = 14) iPSC donors using a reproducible protocol designed to generate three-dimensional neural structures patterned after the cerebral cortex^19, 20^. After 150 days of differentiation, hCOs were exposed to a high-physiological concentration of CLZ for 4-24 weeks to mimic the long-term clinical treatment setting^8^ (Fig. 1a, Methods). The donors were matched on age and sex, 25/28 (89%) were of European ancestry, and one SCZ patient had confirmed TRS based on CLZ use (Suppl. Table 1).

**Figure 1.**
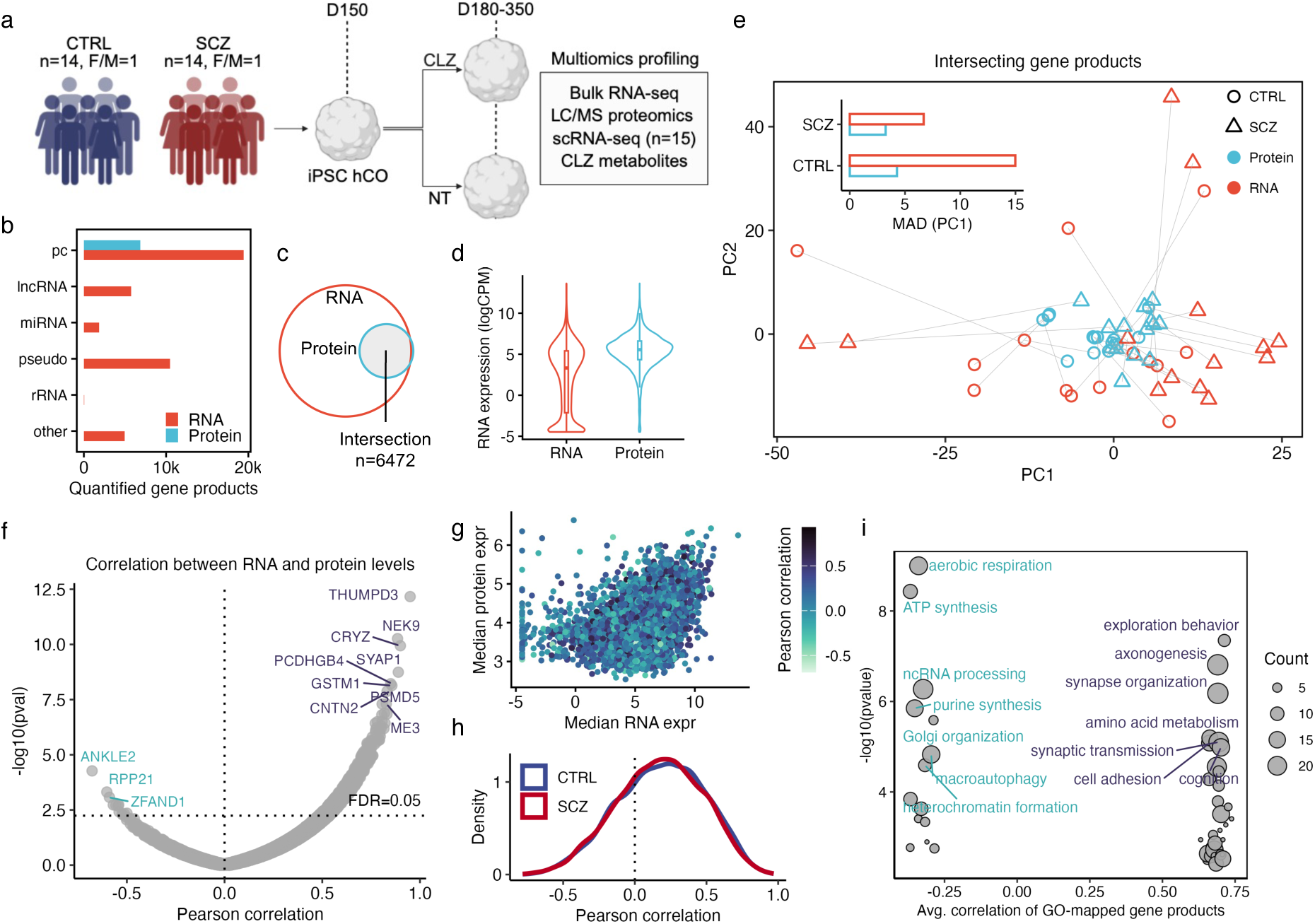
Concordance of transcriptomic and proteomic profiles in hCOs. **a)** Experimental design. CLZ treatment was started at day 150 of differentiation. Bulk RNA-seq, proteomics, measurements of CLZ metabolites were performed at day 180 after four weeks of CLZ exposure. Single-cell RNA-seq (scRNA-seq) was performed at day 350 after 24 weeks of exposure. All concordance analyses were performed on the combined treated and non-treated hCO dataset. F/M: female-to-male ratio. **b)** Number of RNAs and proteins quantified across samples and **c)** the intersection of proteins and protein coding (pc) RNAs. **d)** Violin plot showing the average expression of all RNAs to the left and those intersecting with proteins to the right, indicating that the detected proteins correspond to highly expressed transcripts. **e)** Principal component (PC) plot highlighting the large variability found at the RNA level. The bar plot quantifies this variability by median absolute deviation (MAD) of the first component. **f)** Pearson correlations of the expression levels of the intersecting RNAs and proteins. **g)** Relationship between median RNA expression and protein abundance of intersecting gene products. **h)** The diagnostically stratified distribution of the correlations depicted in f). **i)** Enriched GO terms scored by the average correlation of their mapped gene products.

To characterize the molecular profiles of the derived hCOs in depth, we performed deep bulk RNA-seq and liquid chromatography mass spectrometry (LC/MS) proteomics on 180 days old CLZ-treated and untreated (NT) organoids. We detected and quantified 19,416 protein coding RNAs, 23,101 non-coding RNAs, and 6,849 proteins, of which 6,472 gene products overlapped (Fig. 1b-c). Most of the intersecting proteins corresponded to highly expressed mRNA transcripts (Fig. 1d), suggesting a reduced detection sensitivity of proteomics^21^. Principal component analysis (PCA) based on the intersecting products in the same logarithmic space showed a clear increase in sample variation at the RNA level, with the sample heterogeneity being higher in the CTRL group (Fig. 1e). Molecular studies on brain disorders commonly rely on mRNA expression as a surrogate for protein abundance, but growing evidence suggests limited predictive performance^22, 23^. To assess the relationship between mRNA and protein levels in hCOs, we correlated the abundances of the overlapping gene products using a two-sided Pearson correlation. Correlation coefficients displayed a gene-dependent continuum ranging from -0.69 to 0.93 with a median value of 0.19 (Fig. 1f). Of the significant correlations (FDR < 0.05), 759/770 (98.6%) were positive. The mRNA-protein relationship did not exhibit any dependency on the expression levels of either gene product (Fig. 1g) and was not different between cases and controls (Fig. 1h). To investigate the biological functions in which correlated gene products are involved, we conducted gene ontology (GO) over-representation analysis of the top 5% positively and negatively correlated genes. The over-represented GO terms followed a distinct biological distribution where negatively correlated genes were enriched for basic cellular functions related to energy metabolism and macromolecule processing, while positively correlated genes were enriched for higher brain functions like axonogenesis and synaptic transmission (Fig. 1i). Taken together, these findings corroborate previous evidence showing tighter cellular control of protein translation than mRNA transcription^24^, which is partly due to the high energy expenditure of translation^25^. The results also reinforce previous findings indicating that the strength of transcription control is variable, with more stringent regulation being in place for mRNAs involved in higher brain functions^26^.

### The impact of CLZ on hCO molecular profiles is disease-related and independent of differences in drug metabolism

We next aimed to identify multiomics signatures of SCZ and understand how these are affected by CLZ exposure. Using both the transcriptomic and proteomic data, we performed two sets of differential expression analysis to test for: i) expression differences between SCZ and CTRL hCOs before treatment (baseline differences) and ii) the effects of CLZ treatment by comparing hCOs before and after exposure (treatment effects). The baseline analysis revealed massive changes associated with SCZ with a ∼3-fold increase in the number of differential gene products identified at the RNA-level compared to the protein-level (Fig. 2a). CLZ treatment had a more modest, but robust impact on hCO profiles with increased modulation at the protein-level (Fig. 2a). We found 174 differentially expressed proteins and 63 differential mRNAs that were shared across the two analyses, and their effect sizes (logFCs) displayed a significantly negative correlation (r = -0.32, P = 7.1e-7) at both levels (Fig. 2b). GO analysis of the differential gene products revealed a remarkable enrichment distribution. While downregulated gene products in SCZ hCOs were enriched for synaptic functions, upregulated products were highly enriched for biological processes related to RNA splicing (Fig. 2c). This pattern was reversed by CLZ treatment, which exerted its strongest downregulating effects on proteins involved in RNA splicing. Importantly, this inverse RNA splicing relationship across baseline and treatment comparisons was only detected with the proteomics data, underscoring the importance of an integrative multiomics approach to capture the full repertoire of disease-relevant molecular changes.

**Figure 2.**
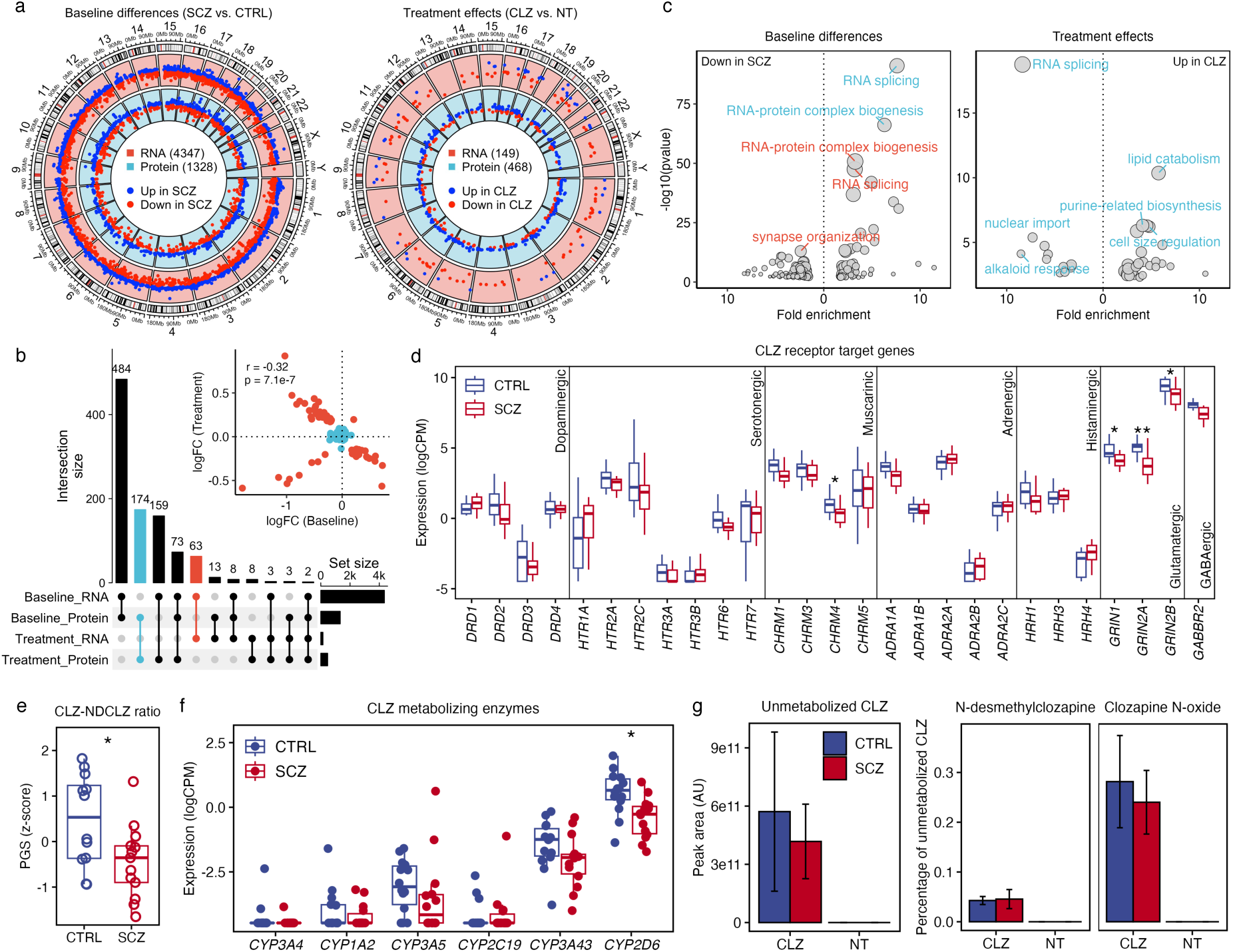
Molecular impact of CLZ on hCO profiles is disease-related and independent of differences in drug metabolism. **a)** Circos plots displaying differentially expressed RNAs and proteins at baseline (SCZ vs. CTRL) to the left and after CLZ treatment (CLZ vs. NT) to the right. **b)** Intersections of differentially expressed gene products at baseline and after treatment. The scatter plot shows a significantly negative correlation of baseline and treatment logarithmic fold changes (logFC) at both RNA and protein levels. **c)** GO enrichment of differential RNAs (red) and proteins (light blue) at baseline (left) and following CLZ treatment (right). **d)** Expression of key CLZ receptor target genes encompassing seven neurotransmitter systems adapted from ref.^9^. **e)** Polygenic score (PGS) of the ratio between CLZ and N-desmethylclozapine (NDCLZ) concentrations (CTRL, n = 12; SCZ, n = 13). **f)** Expression levels of key CLZ metabolizing CYP enzyme genes. **g)** Abundance of unmetabolized CLZ and its two main metabolites in hCO culture media. Metabolite formation is presented as the percentage of remaining CLZ. Non-treated (NT) cultures showed no presence of any compound, indicating the high sensitivity and specificity of the measurements. AU, arbitrary units. **FDR < 0.05, *P < 0.05, two-sided Student t-test.

To investigate whether the effects of CLZ on RNA splicing might be influenced by disease status, we re-ran the treatment analyses in SCZ and CTRL hCOs separately. CLZ had a substantial impact on mRNA expression levels in both cases and controls, but the effect on protein abundance was minimal in SCZ organoids (Suppl. Fig. 1a). Moreover, the effect sizes of the shared gene products (19 mRNAs and 11 proteins) showed no correlation (r = 0.003, P = 0.98), suggesting some dependence of CLZ impact on disease status (Suppl. Fig. 1b-c). Importantly, the downregulation of RNA splicing-related genes was only observed in SCZ organoids, albeit at the RNA-level alone (Suppl. Fig. 1d). While these discrepancies may be partly attributed to the reduced statistical power in the stratified analyses, we investigated other mechanisms that could be involved. CLZ has a highly complex receptor profile^9^, and any major change in receptor availability could influence the cellular action of the drug. We assessed the expression levels of a curated set of 27 key receptor target genes encompassing seven neurotransmitter classes (dopaminergic, serotonergic, muscarinic, adrenergic, histaminergic, glutamatergic, and GABAergic)^9^. Robust expression (logCPM > 0) was observed for most receptor genes, and the muscarinic acetylcholine receptor gene *CHRM4* (encoding the M4 receptor) and the glutamatergic genes *GRIN1*, *GRIN2A*, and *GRIN2B* had significantly reduced baseline expression in SCZ hCOs compared to CTRL (Fig. 2d). This is in line with neuroimaging and postmortem studies demonstrating a widespread decrease in muscarinic M1/M4 receptor levels in SCZ^27^. The G protein-coupled M4 receptor acts through multiple signaling pathways to promote neurogenesis^28^, where alternative splicing plays a crucial role^29^.

In the human body, CLZ undergoes a genetically influenced hepatic metabolism by the CYP system into two main metabolites, N-desmethylclozapine and clozapine N-oxide^30–32^. Using a recent genome-wide association study (GWAS) of CLZ pharmacokinetics (clozapine-desmethylclozapine ratio)^15^, we calculated polygenic scores (PGS) based on genotyping of DNA extracted from iPSC donor blood samples (Methods). We found that the metabolic ratio PGS was significantly lower in SCZ donors (P = 0.022, Fig. 2e). Measurement of the expression levels of key CYP enzymes revealed that SCZ hCOs had reduced expression of *CYP2D6* (Fig. 2f), an enzyme which in the human brain is predominantly expressed in cortical layer III-V neurons^33^. To determine whether these alterations in SCZ organoids translated into actual changes in metabolite concentrations, we quantified the abundance of CLZ, N-desmethylclozapine, and clozapine N-oxide in hCO spent media using ultrahigh-performance liquid chromatography mass spectrometry (UHPLC-MS). All treated hCOs showed low but robust formation of both metabolites (ranging from 0.03-0.52% of unmetabolized CLZ), indicating the presence of CYP enzymatic activity in the organoids (Fig. 2g). Neither N-desmethylclozapine nor clozapine N-oxide were significantly different between cases and controls. In summary, SCZ hCOs were associated with a molecular signature characterized by an upregulation of RNA splicing-related genes, which were reversed by CLZ in a SCZ-specific manner independent of drug metabolic activity, suggesting a convergence between the molecular underpinnings of SCZ and the mechanisms of action of CLZ. Given the weakened statistical power in the stratified comparisons, we concentrated further analysis of treatment effects on the combined SCZ and CTRL data.

### CLZ reverses SCZ-associated transcriptional signatures of intron retention and exon skipping

Alternative splicing is a highly complex and pervasive process taking place in ∼95% of human genes^34^, making it a fundamental regulator of human physiology and disease^35, 36^. It is particularly widespread in the nervous system^29^. The CLZ-induced reversal of the SCZ-associated RNA splicing signature prompted us to elucidate the underlying splicing mechanisms. Leveraging the reference-aligned hCO RNA-seq data, we carried out a comprehensive assessment of multiple alternative splicing events (ASEs) comprising seven major splicing types^37^ (Fig. 3a). We quantified ASEs using the percent spliced in (PSI) metric, representing the ratio between reads including and excluding the alternatively spliced region^38^. After filtering out low-confidence ASEs, we were able to reliably detect and quantify 94,499 unique events across hCO samples, of which 67,134 (71.0%) were intron retention (IR) events and 13,432 (14.2%) were exon skipping (SE) events (Fig. 3b). Differential ASE analysis at baseline identified 636 downregulated and 254 upregulated ASEs (P < 0.05, absolute ΔPSI > 0.05) in SCZ hCOs (Fig. 3c). CLZ exposure resulted in downregulation of 296 and upregulation of 334 events (Fig. 3c). Distributing the differential ASEs across splicing types revealed an over-representation of IR and SE events (Fig. 3d), with the proportion of differential SE events in particular being significantly higher than expected by chance (P < 8.5e-64). Correlating the ΔPSI values of shared IR and SE ASEs between baseline and treatment comparisons showed a negative correlation (r = -0.6, P = 0.0018) of SE events (Fig. 3e), suggesting the ability of CLZ to reverse the regulation of SCZ-associated SE events. We highlight the top reversed SE event in the *KDM2B* gene, which encodes a histone demethylase essential for embryonic development, stem cell regulation, and cellular differentiation, and which is linked to neurodevelopmental disorders^39^. The coverage plot of this event showed a significant decrease in the retention of exon 3 in SCZ hCOs and a complete rescue by CLZ (Fig. 3f). To more closely examine the capability of CLZ to rescue the full SCZ-associated ASE signatures including both shared and unique events, we used connectivity mapping, a computational framework to quantify signature reversal in drug repurposing analyses^40^. We found that CLZ induced a significant reversal of SE, IR, and alternative 3’ splice site (A3SS) signatures (P < 0.0002, Fig. 3g).

**Figure 3.**
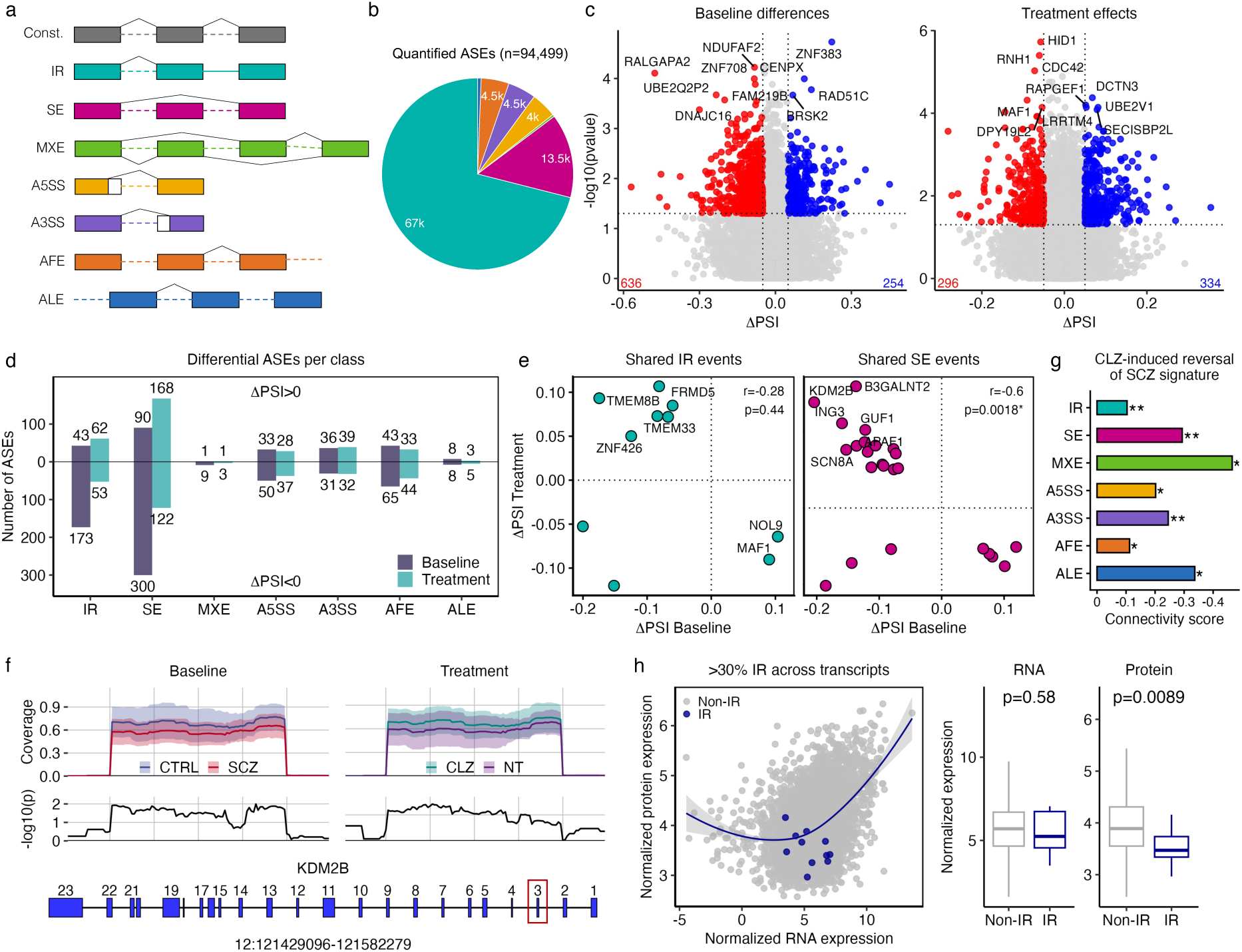
CLZ reverses SCZ-associated transcriptional signatures of intron retention and exon skipping. **a)** Overview of the different alternative splicing event (ASE) types assessed. First row depicts the constitutive isoform. Dotted horizontal line represents skipped intron, while solid line indicates retained intron. IR, intron retention; SE, skipped exon; MXE, mutually exclusive exons; A5SS, A3SS, alternative 5’ and 3’ splice sites; AFE, ALE, alternative first and last exons. **b)** Number of quantified ASEs per type. **c)** Differential ASEs (P < 0.05, absolute ΔPSI > 0.05) identified at baseline (SCZ vs. CTRL) and after CLZ treatment (CLZ vs. NT). A positive ΔPSI value indicates increased inclusion of an event in SCZ/CLZ organoids. **d)** Distribution of up and downregulated ASEs across conditions and splicing types. **e)** Pearson correlation of ΔPSI values for the differential IR and SE events shared between baseline and treatment conditions. **f)** Coverage plot of the top reversed SE event, corresponding to exon 3 in *KDM2B*. **g)** CLZ-induced reversal of splicing-specific SCZ signatures. Connectivity scores ranging from 0 to -1 indicate increasing reversal capacity. **FDR < 0.05, *P < 0.05, weighted Spearman correlation. **h)** Protein levels of most IR genes, defined as genes harboring at least one intron in >30% of transcripts, fall under the regression line, indicating lower abundance than expected based on RNA expression. This shows active degradation of IR transcripts by NMD.

Not all ASEs are expected to be functional, as splicing-induced variation at the mRNA level may not translate into changes at the protein level^35^. If an intron is incorporated into a mature mRNA, a premature termination codon (PTC) may be introduced, leading to the degradation of the mRNA by nonsense-mediated decay (NMD) before the transcript is translated into a protein^41^. IR events thus provide a unique opportunity to examine the extent to which ASEs have functional consequences. We defined IR genes as those harboring one or more introns predicted to be NMD substrates (e.g. due to the presence of an PTC) in at least 30% of the gene’s transcripts. We then compared the normalized mRNA and protein expression levels between IR and non-IR genes in the non-treated hCOs and found that the protein abundance of IR genes was significantly lower than expected based on mRNA levels (Fig. 3h), supporting translational control through NMD-mediated degradation of IR transcripts.

### A cell type-specific alternative splicing program in SCZ

Given the dynamic modulation of alternative splicing across tissues and cell types^42^ and its critical role in governing cell fate during cortical development^43^, we next sought to determine whether the identified alternative splicing alterations exhibited any cell type specificity. We performed ultra-deep single-cell RNA-seq (>80,000 reads per cell) of available hCOs (CTRL: n = 6, SCZ: n = 9) at 350 days of maturation with and without long-term CLZ exposure (24 weeks). Due to reliance on passive surface diffusion of oxygen and nutrients, prolonged organoid maturation may compromise cell viability due to hypoxia and necrotic core formation^44–46^. After stringent quality control to filter out non-viable cells and doublets, we obtained deep single-cell transcriptomics data from 16,274 high-quality cells. We performed shared nearest neighbor (SNN) clustering based on the anchor-integrated data and annotated the clusters using an automated marker-based algorithm^47^ followed by manual refinement. Uniform manifold approximation and projection (UMAP) embedding displayed five progenitor, two neuronal, and one glial population (Fig. 4a-b). Glutamatergic neurons constituted the largest population (48.7%), while *EOMES* expressing intermediate progenitors (IPs, 4.3%) and *ASCL1* expressing early progenitors (EPs, 5.5%) were the least abundant cell types (Fig. 4c). Cell type proportions did not differ across diagnostic and treatment conditions (Suppl. Fig. 2a-b). We used Palantir^48^ to calculate pseudotime based on multiscale space embeddings. Starting from neural precursor cells (NPCs), we identified two differentiation trajectories corresponding to gliogenic and neurogenic cell fates (Suppl. Fig. 2c-f).

**Figure 4.**
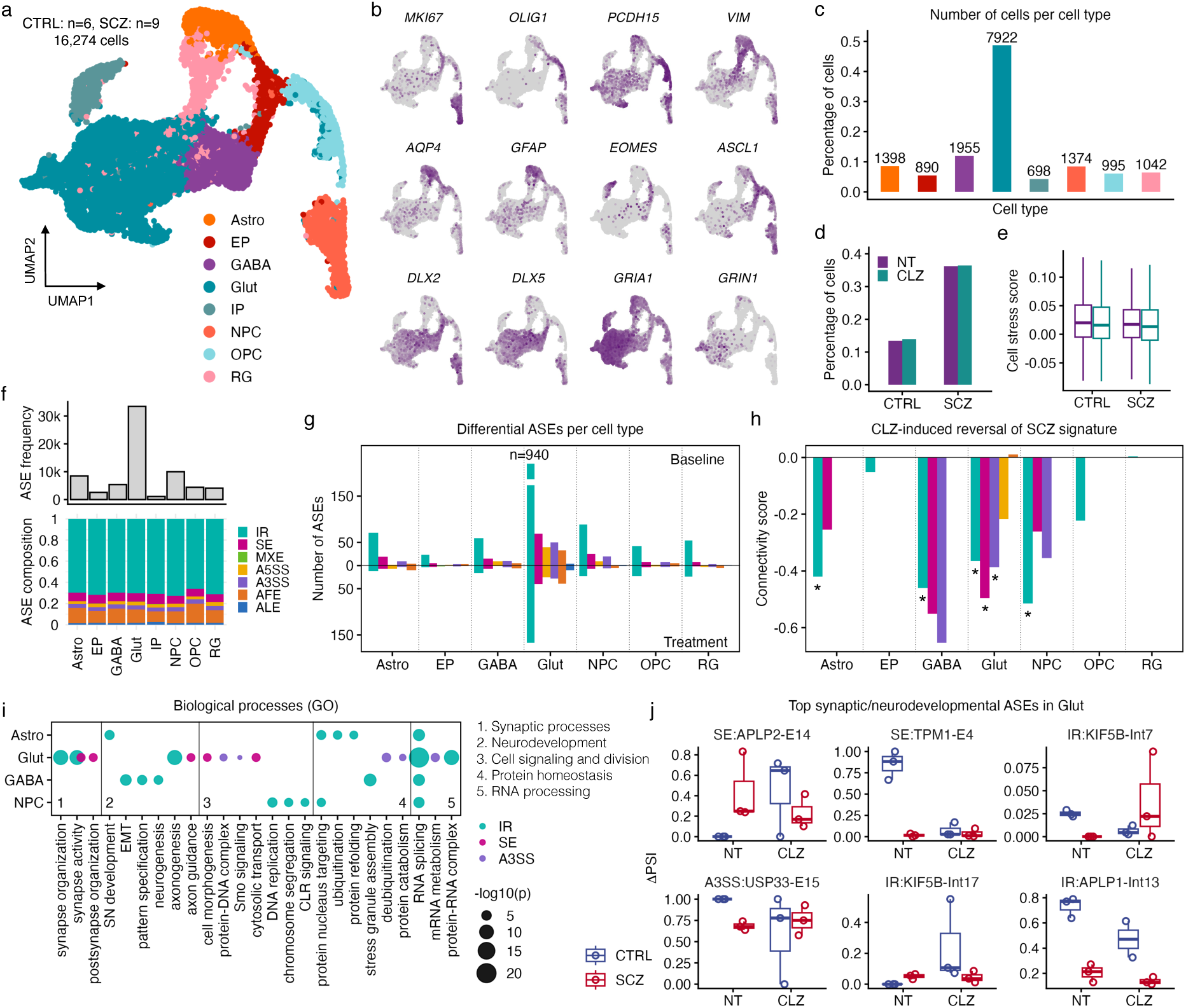
A cell type-specific alternative splicing program in SCZ. **a)** UMAP embedding of the annotated hCO single-cell RNA-seq data. **b)** Marker genes of progenitor, glial, and neuronal cell clusters. **c)** Number of high-quality single cells per cluster. **d)** Proportion of viable cells by diagnosis and treatment condition. **e)** Cell stress scoring showing no significant difference in cellular stress between CLZ-treated and non-treated (NT) hCOs. Paired two-sided Wilcoxon test on donor-level median stress scores. **f)** Frequency and composition of ASEs across cell clusters. **g)** Number of differential ASEs (P < 0.05, absolute ΔPSI > 0.05) at baseline and after CLZ treatment by cell type. No ASEs identified in IPs. **h)** CLZ-induced reversal of splicing-specific SCZ signatures by cell cluster. *FDR < 0.05. **i)** GO enrichment of SCZ-associated splicing events. The five functional GO categories (1-5) range from higher brain functions to basic cellular processes. **j)** Top synaptic and neurodevelopmental ASEs in glutamatergic neurons highlighting the SCZ-specific rescue properties of CLZ. EP, early progenitors; IP, intermediate progenitors; NPC, neural progenitor cells; OPC, oligodendrocyte precursor cells; RG, radial glia.

Our hCOs contained a relatively large proportion of GABAergic neurons (12%) characterized by enriched expression of well-established inhibitory markers like *DLX2* and *DLX5*^49^ (Fig. 4b-c). While the prevailing model of cortical neuron formation stresses the ventral origin of cortical interneurons^50^, a seminal study demonstrated that human cortical progenitors could generate both excitatory and inhibitory neurons^51^. In line with this, varying degrees of ventralization have consistently been observed during dorsal organoid generation^20, 52–55^, reaching up to 15% after 10 months of maturation^56^. The proportion of GABAergic neurons found in our 350 days old hCO cultures is thus well within the expected range. Since high-dose CLZ exposure can have cytotoxic effects, typically manifesting as cell stress and death through metabolic dysfunction and endoplasmic reticulum (ER) stress pathway activation^57^, we assessed the cellular state of our hCOs. We found that the fraction of viable cells was similar across treated and non-treated hCOs in both cases and controls (Fig. 4d), suggesting minimal negative effects of CLZ on cell viability. To determine whether long-term CLZ exposure caused cellular stress, we applied a granular functional filtering method^58^ to calculate cell stress scores based on expression of glycolysis and ER stress marker genes. We found no difference in stress scores between CLZ-treated and non-treated organoids (Fig. 4e). Cells annotated as stressed largely coincided (∼50% overlap) with *VIM* expressing radial glia (RGs), which rely heavily on anaerobic glycolysis for ATP production^59^ (Suppl. Fig. 2g-j). Cell stress scoring after exclusion of RGs confirmed the absence of CLZ-induced cellular stress (Suppl. Fig. 2k).

We next compared the hCO molecular profiles to a recently published single-nucleus transcriptional atlas of the developing human brain^60^, focusing on glial and neuronal cells in the prefrontal cortex (Suppl. Fig. 3a-d). We applied SingleR^61^ to automatically transfer broad cell class (glia, neurons, progenitors) and finer subclass (e.g. astrocytes, Cajal-Retzius, RGs) labels from the reference atlas to hCO clusters based on expression similarity. The transferred class labels were strongly consistent with our assigned labels, with more than 80% concordance for the majority of hCO clusters (Suppl. Fig. 3e-f). Subclass transfer was less concordant for all hCO clusters except for astrocytes, and GABAergic-glutamatergic specification in particular was poorly preserved (Suppl. Fig. 3e-f). These results accord well with previous work reporting poor subtype resolution in organoid clusters compared to primary tissue^55^. We also compared hCO clusters to primary developmental stages spanning the first trimester to adolescence (13 years of age). As expected^62^, we detected strongest matching to the second trimester cortex (Suppl. Fig. 3g). Importantly, this matching was driven by the neuronal cell populations, which are formed earlier in development than glial populations both in vitro and in vivo^17, 60, 63^

To assess the cell type specificity of the identified alternative splicing alterations, we employed a slightly modified version of the bulk pipeline (Methods). The sparse nature of single-cell data (low sequencing coverage and high dropout rates) makes alternative splicing analysis inherently challenging due to reliance on splice junction reads^64, 65^. To overcome this technical limitation, we utilized a synthetic sample generation strategy in which we first pooled together cells of a given cluster from all biological samples within a condition (cases and controls, with and without treatment), and then randomly allocated the cells evenly across three groups, resulting in the construction of 24 synthetic samples per condition (96 samples in total, Suppl. Fig. 4a-d). This strategy, combined with ultra-deep sequencing, enabled us to achieve splice junction coverage rates comparable to those of the bulk data (Suppl. Fig. 4e-f). We detected a total of 68,209 high-confidence ASEs with cell type frequencies mirroring cluster sizes (Fig. 4c,f). The ASE composition was similar across cell types and was dominated by IR events (∼70%, Fig. 4f). Differential ASE analysis by cell type revealed extensive baseline and treatment changes in glutamatergic neurons (Fig. 4g), with a significantly increased proportion of baseline IR events (940 events, P = 3.7e-7). CLZ exposure inverted SCZ-associated IR signatures in astrocytes, GABAergic neurons, glutamatergic neurons, and NPCs (Fig. 4h). In glutamatergic neurons, CLZ additionally reversed SE and A3SS signatures (Fig. 4h). GO analysis further revealed that glutamatergic IR and SE signatures were enriched for genes involved in synaptic and neurodevelopmental processes (Fig. 4i). Closer examination of the top synaptic and neurodevelopmental ASEs in glutamatergic neurons highlighted the reversal and rescue capabilities of CLZ (Fig. 4j).

### Projecting hCO signatures to primary disease profiles and GWAS risk loci

The prenatal-like state of our hCOs raises the question of the extent to which they can recapitulate disease mechanisms and drug effects in the adult brain. To address this question, we projected the hCO disease and treatment signatures to a population-scale single-nucleus atlas of the human prefrontal cortex, representing a diverse range of neurodegenerative and neuropsychiatric disorders^66, 67^. To determine primary disease profiles, we concentrated on patient donors diagnosed with Alzheimer’s disease (AD), Parkinson’s disease (PD), bipolar disorder (BD, type 1 and 2), or SCZ, excluding donors with any comorbidities. We then aggregated the data at class and subclass levels and performed differential expression analyses (Fig. 5a). We then correlated the bulk hCO baseline and treatment signatures to the primary disease profiles. We found that the baseline hCO signatures (i.e. SCZ-associated) exhibited the strongest positive correlation for primary SCZ profiles across cell classes and subclasses (Fig. 5b-c, Suppl. Fig. 5c-d), even stronger than BD, which is closely related to SCZ both genetically and phenotypically^68, 69^. The hCO treatment signatures displayed a remarkable pattern of negative correlation for primary SCZ and BD profiles (Fig. 5b-c), suggesting that our hCO signatures capture real signals in the adult brain. These results were not confounded by age differences across neurodegenerative and neuropsychiatric disorders in the primary atlas (Suppl. Fig. 5e). We also mapped RNA splicing genes (annotated in the GO database) to the primary atlas and found strong enrichment for SCZ in astrocytes, excitatory neurons, and oligodendrocytes (Suppl. Fig. 5f).

**Figure 5.**
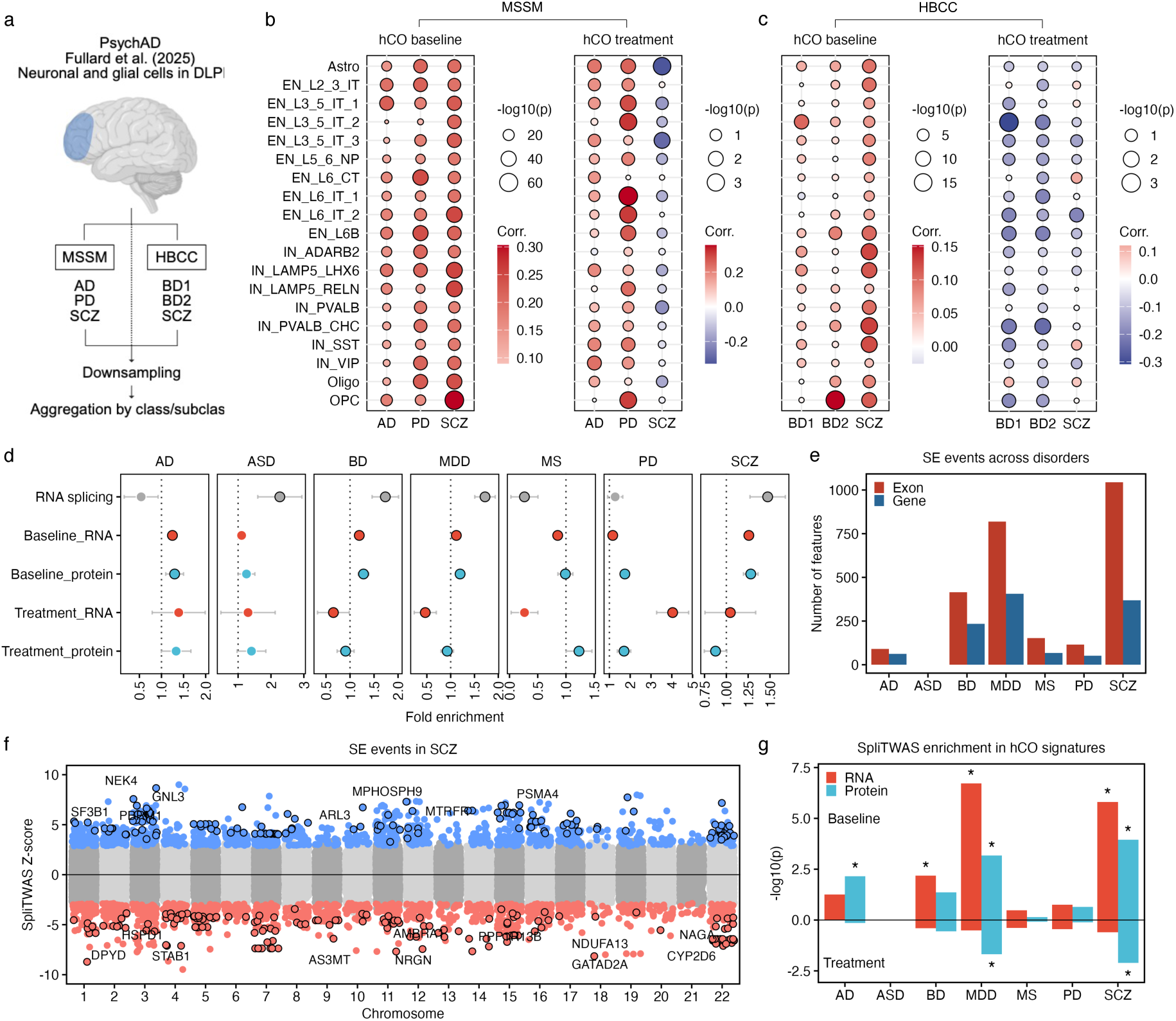
Projecting hCO signatures to primary disease profiles and GWAS risk loci. **a)** Overview of the analysis approach for primary single-nucleus data from the PsychAD consortium^66^. The MSSM and HBCC collections were analyzed separately. Differential expression analyses were conducted on the downsampled and cluster aggregated data. DLPFC, dorsolateral prefrontal cortex. **b,c)** Correlations between baseline and treatment hCO signatures and primary disease profiles in b) MSSM and c) HBCC collections across cell subclasses. EN, IN, excitatory and inhibitory neurons. **d)** Partioning heritability of GWAS traits by the hCO signature gene sets to establish a fold enrichment above the baseline model (dotted vertical line). Dots with black circles indicate a positive delta AIC value above zero, indicating good model fit and reliable enrichment estimate. **e)** Number of SE events with significant SNP heritability tested in SpliTWAS. Red bars indicate the number of affected exons, blue bars the number of unique genes containing these exons. **f)** SpliTWAS associations in SCZ. Colored dots indicate significant associations (FDR < 0.05) and black circles indicate significant colocalization (posterior probability (PP4) > 0.8). **g)** SpliTWAS enrichment of hCO signatures across conditions and GWAS traits. *FDR < 0.05, Fisher exact test.

Finally, we studied whether the hCO signatures display any shared signal with GWAS risk genes. We used a novel functional genomics tool^70^ to partition heritability of a given GWAS trait and determine fold enrichment within genomic regions of interest (ROI), defined by the transcriptional and proteomics hCO signature genes identified in the bulk data as well as the RNA splicing gene set. In addition to the disorders analyzed above, we expanded the GWAS traits to include autism spectrum disorder (ASD), major depressive disorder (MDD), and multiple sclerosis (MS). Enrichment for RNA splicing genes was only seen in neurodevelopmental and neuropsychiatric disorders (Fig. 5d), indicating that RNA splicing mechanisms are less involved in neurological phenotypes. Moreover, bivariate analyses revealed positive genetic correlations (r_g_ > 0, FDR < 0.05) within the defined ROIs across neurodevelopmental and neuropsychiatric disorders (Suppl. Fig. 6). Baseline hCO signatures had highest enrichment for SCZ at both RNA and protein levels (Fig. 5d). While these findings point to an overlap between hCO signatures and SCZ genetic risk, they do not resolve the splicing mechanisms involved. We therefore carried out a splicing-oriented transcriptome-wide association study (SpliTWAS) focused on SE events^71^. We found the largest number of SE events associated with SCZ risk, encompassing 1,044 exons in 369 unique genes (Fig. 5e-f, Suppl. Fig. 7). These events significantly overlapped with both baseline and treatment hCO signatures at the protein level (Fig. 5g). Overall, these results suggest that our organoids may recapitulate some of the genetically determined splicing mechanisms of SCZ and CLZ.

## DISCUSSION

The elucidation of any complex biological process or therapeutical outcome necessitates a multi-faceted approach to thoroughly assess the variety of mechanisms involved. We derived cortical brain organoids from a large case-control cohort of iPSC lines and employed a comprehensive multiomics strategy to decipher the long-term cellular mechanisms of action of CLZ in human brain cells. Transcriptomics and proteomics profiling revealed an RNA splicing signature in SCZ organoids which was reversed by CLZ. While this effect was more pronounced in SCZ organoids, it was not driven by differences in the rate of CLZ metabolism. In-depth bulk and single-cell analyses uncovered a high-resolution alternative splicing program in SCZ, emphasizing the role of exon skipping and intron retention in glutamatergic neurons. This druggable program, which was found to target genes and pathways involved in synaptic and neurodevelopmental processes, recapitulated splicing-associated disease mechanisms in primary human tissue and captured genetic signals of SCZ risk. Importantly, prolonged CLZ exposure did not negatively impact cell viability and stress, indicating minimal confounding by adverse drug effects.

Among all human tissues, the brain exhibits the most extensive patterns of alternative splicing and the highest degree of isoform diversity^72^, features that both support higher-order human cognitive functions and contribute to neuropsychiatric disease risk^73^. SCZ risk is strongly influenced by splicing variants and isoform dysregulation^74–77^, particularly in excitatory neurons during the first two trimesters of development^73, 78, 79^. These observations align well with our finding of convergence between hCO signatures and primary disease mechanisms and GWAS signals, and suggest that the splicing effects captured by our organoids represent the early-stage impact of SCZ risk variants. Novel therapeutics targeting aberrant splicing has proven successful for the treatment of a variety of diseases, ranging from spinal muscular atrophy to cancer^36^. These therapeutic agents often act by recognizing specific RNA splicing regulatory elements to modulate splicing, often by blocking abnormal recruitment of splicing factors^80^. The reversal of SCZ-related IR and SE events by CLZ in our hCOs suggests that it may act by a similar mechanism upstream of transcription. While these results are encouraging, it remains unclear whether the effects on splicing fully explain the unique clinical efficacy of CLZ or whether these actions are shared by other antipsychotics. Previous work indicates that olanzapine and haloperidol may also act through splicing regulation^81, 82^, but these studies were conducted in mice and did not report the strong SCZ-associated splicing reversal capability observed here.

Our hCO cohort included only one patient with confirmed TRS, raising the question of how relevant the identified alternative splicing program is to the underlying mechanisms of TRS and CLZ efficacy. This question hinges on the extent to which TRS represents an exacerbated manifestation of SCZ or an orthogonally distinct phenotype. Some studies report no common genetic overlap between SCZ and TRS^83, 84^, but others have found significant associations between TRS status and polygenic risk for SCZ^4, 84–87^. The proportion of rare SCZ variants has also been found to be enriched in TRS cases compared to non-TRS cases^87^, suggesting that the biological underpinnings of TRS at least partly overlap with those of SCZ. Our hCO patient donors had a high load of SCZ risk variants (Suppl. Table 1), and we found that the CLZ-induced reversal of the splicing signature was more pronounced in SCZ hCOs, both of which speak to the relevance of the identified splicing program to TRS and CLZ response. Nevertheless, the inclusion of both CLZ-responsive and non-responsive TRS patient groups will be needed to confidently resolve the therapeutic mechanisms of action of CLZ.

An innovative aspect of the present study was the thorough assessment of CYP-mediated CLZ metabolism. In the human body, CLZ is extensively (∼80%) metabolized in the liver through demethylation and oxidation to form N-desmethylclozapine and clozapine N-oxide, only the former of which exhibit some psychotropic activity^88^. CLZ metabolism is primarily carried out by CYP1A2 and CYP3A4/5 enzymes, while CYP2D6 plays a minor role^88^. Our SCZ donors had a reduced metabolic rate PGS, but this difference did not translate into reduced expression of *CYP1A2* and *CYP3A4/5* nor an actual reduction in CLZ metabolite concentrations in the organoids. These results are in accordance with previous work suggesting a modest impact of polygenic load on enzymatic activity, explaining less than 10% of variance in CLZ pharmacokinetics^15^. Of the CYP enzymes analyzed, *CYP2D6* had the highest expression in hCOs and was significantly downregulated in SCZ organoids. This finding was corroborated by the SpliTWAS analysis, which found that splicing changes in *CYP2D6* were associated with SCZ susceptibility. A recent study provided further support for a role of *CYP2D6* splicing in SCZ^89^. Importantly, *CYP2D6* expression was found to be particularly high prenatally in the frontal cortex^89^. In addition to drug metabolism, *CYP2D6* plays an important role in the biotransformation and neurotransmission of central dopamine and serotonin^90^, which may explain its link to SCZ pathology.

In conclusion, our comprehensive investigation into the long-term molecular effects of CLZ in human brain cells identified a druggable alternative splicing program in SCZ. This encourages increased efforts to understand the intricate splicing mechanisms involved, which may open exciting avenues for the development of a new generation of antipsychotics.

## ACKNOWLEDGEMENTS

This work was supported by the Research Council of Norway (223273, 248828, 300309, 334920), the South-Eastern Norway Health Authorities (2022087), and the EU Horizon Psych-STRATA project (101057454). We thank Sarah Åsheim, Elin Inderhaug, Asbjørn Holmgren, and Denis Reis de Assis for their excellent technical assistance. We also want to extend our thanks to the Norwegian Core Facility for Human Pluripotent Stem Cells Research Centre for the reprogramming of fibroblasts and the use of the BSL-2 laboratory. RNA sequencing and the preprocessing pipeline were provided by the Norwegian Sequencing Centre (www.sequencing.uio.no), a national technology platform hosted by the University of Oslo and supported by the Functional Genomics and Infrastructure programs of the Research Council of Norway and the South-Eastern Regional Health Authorities.Mass spectrometry-based proteomic analyses were performed by the Proteomics Core Facility, Department of Immunology, University of Oslo and Oslo University Hospital, which is supported by the Core Facilities program of the South-Eastern Norway Regional Health Authorities. This core facility is also a member of the National Network of Advanced Proteomics Infrastructure (NAPI), which is funded by the Research Council of Norway INFRASTRUKTUR program (295910).

## AUTHOR CONTRIBUTIONS

S.D., K.O., O.A.A., V.M.S. and A.T. provided funding for this work. S.D. and J.R.O. conceived, designed and supervised the work. I.A.A. carried out bioinformatic analyses and writing. J.R.O. performed stem cell experiments. S.D. and O.A.A. were responsible for iPSC donor recruitment and sample collection. E.M. carried out UHPLC-MS analysis of CLZ metabolites. N.P. performed bivariate GSA-MiXeR analyses. E.K. calculated metabolic rate polygenic risk scores. All authors provided critical feedback on the manuscript.

## METHODS

### Collection and characterization of human biospecimens

Skin biopsy donors were recruited through the Norwegian Thematically Organized Psychosis (TOP) study^91, 92^. Fibroblasts were isolated from 14 CTRL individuals and 14 SCZ patients as previously described^17, 18^. Cases and controls were matched on age and sex, and all except for three donors were of white European ethnicity. One patient had confirmed TRS based on CLZ use (Suppl. Table 1). For genotyping and PGS calculation, DNA was extracted from blood samples collected at inclusion in the TOP study. As previously reported^93^, genotyping was performed on the Illumina Human Omni Express-24 v.1.1 BeadChip, and standard pre-imputation quality control was performed using PLINK 1.9^94^. A PGS of CLZ metabolism was calculated using PRS-CS^95^ with default parameters based on a recent GWAS of clozapine-to-N-desmethylclozapine metabolic ratio^15^.

### Reprogramming and iPSC characterization

Skin fibroblast from the 28 donors were grown and reprogrammed to iPSCs with Sendai virus using the CytoTune iPS 2.0 Sendai Reprogramming Kit (Thermo Fisher Scientific, A16517) as previously described^96^. Each iPSC line was subjected to rigorous quality control and characterization by phenotyping, regular monitoring of morphology and pluripotency marker expressions at the Norwegian Core Facility for Human Pluripotent Stem Cell Research Centre.

### Generation of hCOs from iPSCs

Donor iPSCs were differentiated to cortical organoids (hCOs) following an established protocol with proven reproducibility^19, 20^. The iPSCs were first dissociated into single cells by incubation with Acutase (Sigma, A6964) at 37°C for 5 min. Aggrewell 800 (STEMCELL Technologies, 34815) was used to obtain homogeneously sized hCOs. To prevent cell adhesion, plates were pre-treated with anti-adherent rinsing solution (STEMCELL Technologies, 07010) following manufacturer’s instructions. Approximately 18×10^6^ cells in Essential 8 Flex medium supplemented with the ROCK inhibitor Y-27632 (10µM, Miltenyi Biotec) were added per well of a 6-well Aggrewell 800, centrifuged for 3 min at 100g, and incubated at 37 °C with 5% CO2. After 24 hours of incubation, corresponding to day 0 of differentiation, cell aggregates were collected from each microwell by pipetting up and down with a P1000 tip (with the end cut) and transferred to ultra-low-attachment plastic dishes (Thermo Fisher Scientific, 15297905) in Essential 6 medium (Life Technologies, A1516401) supplemented with dual SMAD inhibitors, dorsomorphin (2.5 μM, Sigma-Aldrich, P5499) and SB431542 (10 μM, Tocris, 1614), together with Wnt inhibitor XAV939 (2.5 μM, Tocris, 3748). Medium was changed daily from day 2 to day 6 (E6 plus inhibitors). After day 6, suspended hCOs were transferred to neural medium containing Neurobasal-A (Life Technologies, 10888), GlutaMax (1:100, Life Technologies, 35050) and B-27 supplement without vitamin A (Life Technologies, 12587). Neural medium was supplemented for 19 days with 20 ng/ml epidermal growth factor (EGF, R&D Systems, 236-EG) and 20 ng/ml fibroblast growth factor (FGF, R&D Systems, 233-FB), with medium changed daily in the first 10 days and every other day for the following 9 days. From day 24 until day 43, neural medium was supplemented with 20 ng/ml brain-derived neurotrophic factor (BDNF, Peprotech, 450-02) and 20 ng/ml neurotrophin 3 (NT3, Peprotech, 450-03) to promote neural differentiation, with medium changed every other day. From day 43 onwards, only neural medium without growth factors was changed every 3-4 days.

### Clozapine administration

At day 150 of differentiation, hCOs were exposed to a supraphysiological concentration of 10 µM CLZ (Sigma, C6305) diluted in DMSO, which was used as vehicle control. High-physiological dosages are routinely used in in vitro studies^97, 98^, especially in neural cultures to evoke maximal electrical activity^99^. This is particularly important in hCO tissues given their dense 3D structure and inherent diffusion barriers. In a clinical context, long-term CLZ treatment of up to 6 weeks is required to achieve 30% clinical response rate^8, 100^. By 12 weeks of treatment, 40-50% of patients respond adequately, providing a lower threshold for the recommended treatment duration^8, 100^. To recapitulate this clinical long-duration setting in vitro, hCOs were exposed to CLZ for one month from day 150-180 of differentiation, with medium and drug refreshment every 3-4 days. At day 180, one hCO per donor line was processed for bulk RNA-seq and one was processed for LC-MS/MS proteomics, while the remaining hCOs were kept in culture with medium and drug refreshment every 3-4 days until day 350 of differentiation.

### Bulk RNA-seq processing

Total RNA was extracted from CLZ-treated and non-treated hCOs from all 28 donors (n=56 samples in total) using the RNeasy Plus Mini Kit (Qiagen). RNA yield was quantified with a NanoDrop 8000 Spectrophotometer (NanoDrop Technologies) and RNA integrity was assessed with Bioanalyzer 2100 RNA 6000 Nano Kit (Agilent Technologies). Library preparation and paired-end RNA-sequencing were carried out at the Norwegian High-Throughput Sequencing Centre (www.sequencing.uio.no). Briefly, libraries were prepared with the TruSeq Stranded mRNA kit from Illumina which involves poly-A purification to capture coding as well as several non-coding RNAs. The prepared samples were then sequenced in one batch on an Illumina NovaSeqX platform at an average depth of 74 million reads per sample, using a paired-end read length of 150 bp. Raw sequencing reads were quality assessed with FastQC (Babraham Institute). To pass the initial QC check, the average Phred score of each base position across all reads had to be at least 30 (99.9% base calling accuracy). Reads were further processed by cutting individual low-quality bases and removing adapter and other Illumina-specific sequences with Trimmomatic V0.32^101^. Any remaining phiX and Sendai virus sequences were filtered out with BBTools^102^. The splice-aware alignment tool HISAT2^103^ was then used to first build a transcriptome index based on Ensembl annotations, and then to map the clean reads to the human GRCh38 reference genome. To quantify gene expression levels, mapped reads were summarized at the gene level using featureCounts^104^ guided by Ensembl annotations.

### Protein extraction and sample preparation

Proteins were extracted from all 56 hCO samples by cell lysis using RIPA buffer (Sigma, R0278) with protease and phosphatase inhibitor cocktail (Thermo Fisher Scientific, 1861281) and 5 mM EDTA (Thermo Fisher Scientific, 1861274). Total protein concentration was measured with BCA protein assay kit (Thermo Fisher Scientific, 23225). From each lysate, 10 µg of protein was precipitated with 70% acetonitrile on magnetic beads (MagReSyn Amine, Resyn Biosciences). The proteins were washed on the beads with 100% acetonitrile and 70% ethanol and then resuspended in 50 µl 50 mM ammonium bicarbonate containing 10 mM DTT for reduction of cysteines. Samples were then incubated at 37°C for 40 minutes. To alkylate the proteins, 50 µl of 30 mM IAA in 50 mM ammonium bicarbonate was added to the samples, which were incubated at room temperature in the dark for 30 minutes. Trypsin (0.5 µg) was added to each sample for on-bead protein digestion at 37°C overnight. The resulting peptides were concentrated and desalted on Evotip (Evosep Biosystems, Denmark) for mass spectrometry analysis.

### LC-MS/MS proteomics

LC-MS/MS analysis was carried out with the Evosep One LC system (EVOSEP Biosystems, Denmark) coupled to a timsTOF Pro2 mass spectrometer, using a CaptiveSpray nano electrospray ion source (Bruker Corporation, Germany). Digested peptides (4 ul) were loaded onto a capillary C18 column (Evosep Biosystems, Denmark). Peptides were separated at 40°C using the standard 30 sample/day method from Evosep. The timsTOF Pro2 mass spectrometer was operated in DIA-PASEF mode. Mass spectra for MS were recorded between 100-1700 m/z. Ion mobility resolution was set to 0.85–1.30 V·s/cm over a ramp time of 100 ms. The MS/MS mass range was limited to m/z 475-1000 and ion mobility resolution to 0.85-1.27 V s/cm to exclude singly changed ions. The estimated cycle time was 0.95 seconds with 8 cycles using DIA windows of 25 Da. Collisional energy was ramped from 20eV at 0.60 V s/cm to 59eV at 1.60 V s/cm. Raw data files from LC-MS/MS analyses were submitted to DIA-NN (v1.8.1)^105^ for protein identification and label-free quantification using the library-free function. Parameters were as follows: carbamidomethyl (C) was set as a fixed modification; the trypsin without proline restriction enzyme option was used, with one allowed miscleavage; peptide length range was set to 7-30 amino acids; mass accuracy was set to 15 ppm; and the FDR allowed was 0.01. The UniProt human database (May 2024) was used for the database searches. The DIA-NN output data was loaded into Perseus (v1.2.11.0)^106^ for quality evaluation. Identifications from potential contaminants and reversed sequences were removed, and normalized intensities (LFQ) were log10-transformed. A requirement of at least 70% valid values in at least one group was applied to filter the results. All zero intensity values were replaced using noise values of the normal distribution of each sample. The log10-transformed normalized intensities were used in all statistical analyses.

### Differential expression

Differential expression analysis of both bulk RNA-seq and proteomics data was carried out with limma-voom^107^. Gene counts were first filtered using the filtreByExpr function in the edgeR package^108^, and only protein coding and long non-coding RNAs, the two most abundant RNA species, were retained. Counts were then normalized using trimmed mean of *M* values^109^ as implemented in the calcNormFactors function from edgeR^108^. To minimize the potential effects of outlier samples, we leveraged a strategy using quantitative quality weights^110^ as implemented in the voomWithQualityWeights function from limma^107^. Sex and age were included as model covariates in all baseline analyses (i.e. SCZ vs. CTRL). Genes with FDR < 0.1 were considered differentially expressed. GO enrichment tests of differentially expressed genes were conducted with the over-representation analysis tool clusterProfiler^111^ using the enrichGO function, selecting BP (biological processes) as the ontology of interest. Term redundance was identified and reduced by calculating semantic similarity^112^. GO terms with FDR < 0.1 were considered significantly enriched.

### UHPLC-MS of CLZ metabolites

To detect and quantify CLZ and its two main metabolites (clozapine N-oxide and N-desmethylclozapine), hCO spent media samples were submitted to a therapeutic drugmonitoring (TDM) service at Center for Psychopharmacology in Oslo, Norway. Samples were prepared by protein precipitation in a semiautomated manner using a Microlab Star pipetting robot (Hamilton) and analyzed using a vanquish ultrahigh-performance liquid chromatography (UHPLC) system (Thermo Fisher Scientific). Chromatographic separation was obtained using an XBridge BEH C18-column (2 *μ*m, 2.1 × 75 mm, Waters) with a gradient elution consisting of ammonium acetate buffer (pH = 4.8) and acetonitrile (20%–52%) at 35°C. The UHPLC system was coupled with a high-resolution mass spectrometer equipped with an Orbitrap detector set to positive ionization mode, scanning data at a resolution of 70,000 across a mass range of 100–1500 Da, allowing precise mass separation up to the fourth decimal. Quantification of CLZ was conducted using a standard calibration curve, with a lower detection limit of 20 nmol/L. Identification and quantification of metabolites were based on accurate mass (with a mass tolerance of 5 ppm for the protonated molecular ion), isotope ratio, and evaluation of the MS/MS spectra consistent with chemical structures of CLZ-derived compounds. Semiquantification of metabolites was determined in arbitrary units based on chromatographic peak intensities with detector responses assumed to be linear and similar to that of CLZ. The metabolite-to-CLZ levels (metabolic ratios) were used as a measure of metabolite formation.

### Bulk alternative splicing analysis

Bulk RNA-seq alignment files (BAM files) were processed and collated with the R package SpliceWiz^37^. ASEs were quantified using the percent spliced in (PSI) metric, representing the ratio of mRNA transcripts that include the alternatively spliced region versus those that exclude it. Quantification of IR events followed the IRFinder^113^ approach, using the 30% trimmed mean depth of coverage across the measured intron. Abundance of the remaining ASEs (SE, MXE, A5SS, A3SS, AFE, ALE) was estimated using junction read counts, which do not require length normalization. Alternative splicing annotations were based on the human GRCh38 reference genome. Low-confidence ASEs were excluded using default filters. Differential ASE analyses were conducted on the PSI values after replacing missing values with the median across samples and excluding events with >80% non-unique values. Mann-Whitney test was used to identify baseline differences, while Wilcoxon Signed-Rank test was used for treatment effects. ASEs with P < 0.05 and absolute ΔPSI > 0.05 were considered differentially expressed. The connectivityScore function as implemented in the PharmacoGx^114^ package was used to calculate connectivity scores of signature reversal based on a weighted Spearman statistic (GWC).

### Dissociation of hCOs and sample preparation

Each hCO at differentiation day 350 was gently cut into small pieces and transferred to a 1.5 mL Eppendorf tube for digestion in 1 mL Trypsin-EDTA (0.05%) solution (Thermo Fisher Scientific, 25300054). Samples were incubated at 37°C for 10 minutes. Digestion was terminated by transferring the cells to a 15 ml tube containing 3 ml of neural medium. The cell suspensions were then centrifuged for 10 minutes at 300 g, resuspended in 1 ml neural medium, and filtered through a 40 µm cell strainer to reduce aggregation. Cell counting and measurement of cell viability were performed on a NucleoCounter NC-200 (ChemoMetec). The cells were fixed and permeabilized using the Parse Biosciences Evercode Nuclei Fixation Kit v3 (Parse Biosciences). Split-pool combinatorial barcoding and library preparation were carried out using the Evercode WT Mega v3 kit (Parse Biosciences) according to manufacturer’s instructions.

### Single-cell RNA-seq and data processing

Libraries were sequenced at the Norwegian Sequencing Center on an Illumina NovaSeq X flow cell using 100 bp paired-end sequencing with 15% PhiX. Approximately 80,000 pair-end reads were generated per cell. FASTQ files for each sublibrary (15 in total) were run through the Parse Biosciences split-pipe (v1.5.1) pipeline for barcode correction, trimming of reads, and alignment to the human GRCh38 reference genome. Automatic filters (cell size distribution, mitochondrial content, genes vs. transcripts, and doublets) were applied and the resulting count matrices (‘DGE_filtered’) were imported to Seurat (v5.0.0). Doublets were identified with scDblFinder^115^ and singlet barcodes with less than 10% mitochondrial reads and with 500-10,000 features were retained. The filtered data was processed following standard best practices for Seurat v5, including log-normalization of gene counts and anchor-based CCA sample integration. Clusters were annotated through a two-step procedure. First, ScType^47^ was used for an initial automatic annotation of brain cell types. Second, each cluster annotation was manually refined based on differential expression of marker genes and literature searches. For pseudotime analysis, the Palantir algorithm^48^ as implemented in SeuratExtend^116^ was used to first generate diffusion map and multiscale space embeddings based on CCA reductions. These embeddings were then used to calculate pseudotime to determine the developmental trajectory and cell fate decisions from a user-defined starting cell. To quantify cell stress, a granular functional filtering method^58^ was applied, which calculates a functional score per granule (small group of cells with similar expression profiles) based on the expression of glycolysis and ER stress genes as positive filters, and the expression of gliogenesis and neurogenesis-related genes as negative filters. Due to confounding with the RG population, all cells were retained in downstream analyses, whether marked as stressed or not.

### Downstream data analysis

Due to the inherent correlation between cells originating from the same sample, the AggregateExpression function from Seurat v5 was used to create summed counts (‘pseudobulk’) for each cell identity per sample. Differential expression analyses were then carried out on these counts following the bulk procedure described above. The sparse nature of single-cell data (high dropout rates and low coverage) makes alternative splicing analysis technically challenging due to its reliance on splice junction reads. To overcome this technical limitation, we applied a supervised synthetic sample generation strategy. For each condition (cases and controls, with and without treatment), cells of a given identity from all biological samples were first pooled and then randomly, but evenly, allocated across three groups, resulting in the construction of 24 synthetic samples per condition (96 in total). The unique cell IDs from each synthetic sample were used to generate sample-specific BAM files. ASE detection and quantification followed the same SpliceWiz^37^ pipeline as with the bulk RNA-seq data, except for the filtering of low-confidence events, which were slightly loosened to increase detection rates (filters passed in >70% of samples, 35% IR coverage, and 20% non-IR coverage). The edgeR wrapper function ASE_edgeR was used to test for differential ASEs, excluding the IP cell population due to low read counts. Events with P < 0.05 and absolute ΔPSI < 0.05 were considered differentially spliced.

### Cell label transfer from the human developing brain single-nucleus atlas

Processed data of a recent primary developing brain atlas^60^ were downloaded as an AnnData object from CELLxGENE (https://cellxgene.cziscience.com/collections/ad2149fc-19c5-41de-8cfe-44710fbada73). Non-neural cells, cells annotated as unknown, and cells originating from the primary visual cortex (V1) were excluded. Only protein coding genes were retained. The data were downsampled to 20,000 randomly selected cells to reduce computational requirements. To compare our hCO single-cell clusters to their primary counterparts, we used SingleR^61^ to automatically transfer cell labels from the reference atlas to hCO clusters based on expression similarity. To speed up the process, we applied the aggr.ref option to train the SingleR classifier on aggregate reference clusters rather than each cell individually.

### Projection of hCO signatures to primary disease profiles

To assess the relationship between our hCO signatures (disease and treatment) and disease profiles from primary brain tissue, we obtained snRNA-seq data from the PsychAD Consortium^66^, representing a population-scale cross-disorder atlas of the human prefrontal cortex. Raw count data were downloaded as AnnData objects from CELLxGENE (https://cellxgene.cziscience.com/collections/84ce6837-548d-4a1f-919f-0bc0d9a3952f). The MSSM and HBCC cohorts displayed biobank-related batch effects and were processed and analyzed separately (Suppl. Fig. 5a-b). Healthy controls and patient donors with a diagnosis of either AD, BD, SCZ, or PD and no comorbidities were included. Only cells of the neuronal and glial lineages (Astro, EN, IN, OPC, and Oligo) and protein coding genes were retained for analysis. To identify disorder-associated genes, differential expression analyses were performed separately in each cell class and subclass. The data were downsampled to 10,000 random cells and the counts were collapsed by donor using the collapse_counts function from presto^117^. The limma-voom^107^ workflow, including gene filtering, normalization, and quantitative quality weighting, was applied. The statistical models were adjusted for donor age, sex, genetic ancestry, and the number of genes detected. For each gene, a significance score was calculated as: score = -log10(P)*sign(logFC). The Pearson correlation of the significance scores from organoid and primary data was used to quantify concordance.

### Bivariate GSA-MiXeR analysis

Bivariate GSA-MiXeR, a novel functional genomics tool^70^, was used to partition the heritability of GWAS traits within genomic regions of interest (ROI), defined by the hCO signature genes. Both trait-specific fold enrichment of heritability (univariate) and genetic correlation between two traits within signature ROI (bivariate) analyses were performed. As input, we used the differentially expressed RNA and protein sets from the bulk hCO analyses and summary statistics from the latest Psychiatric Genetics Consortium GWAS of AD^118^, ASD^119^, BD^120^, MDD^121^, and SCZ^122^. In addition, we used summary statistics for PD from the GWAS by Nalls et al. (2019)^123^ and for MS from the International Multiple Sclerosis Genetics Consortium^124^. All GWAS data included participants of European ancestry only. In line with the original GSA-MiXeR model^125^, fold enrichment values have an associated standard error and delta AIC value but no p-values. We considered an ROI as enriched when the fold enrichment value was above one and the delta AIC > 0, indicating a good model fit and reliable enrichment estimate.

### Splicing-specific TWAS

Splicing-specific transcriptome-wide association studies (TWAS) were carried out with SpliTWAS^71^, an exon skipping-oriented implementation of the FUSION^126^ framework. Precomputed PSI weights of exon skipping from the PsychENCODE BrainGVEX^127^ dataset available at https://zenodo.org/records/10015110 were used. This dataset was derived from postmortem dorsolateral prefrontal cortex samples from 344 individuals (91 cases and 253 controls) of European ancestry. A total of 12,609 exons (in 5,978 genes) with significant *cis*-SNP heritability were tested. LD information was obtained from the 1000 Genomes non-Finnish European reference. Rare autosomal SNPs with MAF < 0.01 were filtered out with PLINK2^94^. The same GWAS summary statistics as above were utilized. We also ran colocalization^128^ tests on any splicing-trait associations with a SpliTWAS p-value less than 0.05 (--coloc_P 0.05 flag in FUSION). Splicing-trait associations with an FDR < 0.05 were considered significant, and associations with a COLOC posterior probability (PP4) > 0.8 were considered colocalized.

### Ethical compliance

Informed consent was obtained from all participants, and the study was approved by the Norwegian Data Protection Agency and the Regional Ethics Committee of the South-Eastern Norway Regional Health Authorities (REK grant: 2012/2204). The authors declare that all procedures contributing to this work comply with the ethical standards of relevant guidelines and regulations.

