## Supplementary Information for "Integrative multiomics profiling of cortical brain organoids reveals a druggable alternative splicing program in schizophrenia"

**Suppl. Table 1. Clinical and demographic characteristics of hCO donors**

| Characteristic | CTRL, n = 14 | SCZ, n = 14 | Test statistic | P value |
|---|---|---|---|---|
| Subdiagnosis | | | | |
| Schizophrenia, n (%) | | 10 (71.4) | | |
| Schizoaffective, n (%) | | 3 (21.4) | | |
| Schizophreniform, n (%) | | 1 (7.2) | | |
| TRS status | | | | |
| TRS | | 1 (7.2) | | |
| non-TRS | | 13 (92.8) | | |
| Age | | | t = 0.77 | 0.45 |
| Mean (SD) | 33.1 (10.6) | 29.9 (11.5) | | |
| Median (range) | 30.5 (18-56) | 25.0 (19-51) | | |
| Sex | | | $\chi 2 = 0$ | 1 |
| Male, n (%) | 7 (50) | 7 (50) | | |
| Female, n (%) | 7 (50) | 7 (50) | | |
| Ethnicity | | | $\chi 2 = 3.36$ | 0.19 |
| European, n (%) | 14 (100) | 11 (78.6) | | |
| African, n (%) | | 1 (7.1) | | |
| Mixed, n (%) | | 2 (14.3) | | |
| Polygenic risk score SCZ | | | t = -2.50 | 0.026* |
| Mean (SD) | -0.83 (1.52) | 0.33 (0.5) | | |
| Median (range) | -1.04 (-3.1-1.5) | 0.28 (-0.4-1.4) | | |
| Medication | | | | |
| Antipsychotics, n (%) | | 11 (78.6) | | |
| Antidepressants, n (%) | | 2 (14.3) | | |
| PANSS | | | | |
| Positive symptoms, mean (SD) | | 13 (4.2) | | |
| Negative symptoms, mean (SD) | | 15 (5.6) | | |
| General psychopathology, mean (SD) | | 32 (9.1) | | |
| Total score, mean (SD) | | 59 (13.9) | | |

TRS, treatment-resistant SCZ based on CLZ prescription; PANSS, Positive and Negative Symptom Scale. *P < 0.05, two-sided Student t-test.

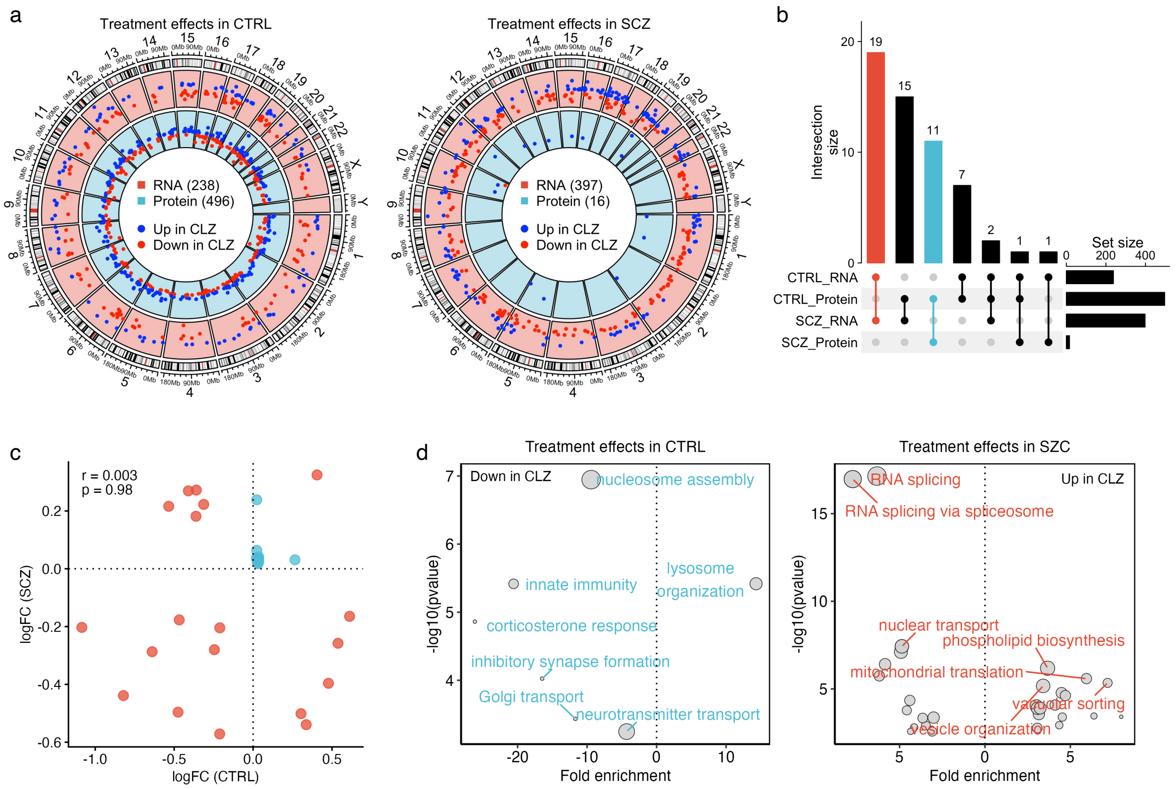


**Suppl. Figure 1. Molecular impact of CLZ on hCO profiles in CTRL and SCZ organoids. a)** Circos plots displaying differentially expressed RNAs and proteins in CTRL (left) and SCZ (right) organoids. **b)** Intersections of differentially expressed gene products across CTRL and SCZ. **c)** Dot plot showing no correlation between CTRL and SCZ logarithmic fold changes (logFC) at RNA or protein levels, indicating disease-dependent effects of CLZ. **c)** GO enrichment of differential gene products in CTRL (left) and SCZ (right) hCOs after CLZ exposure.

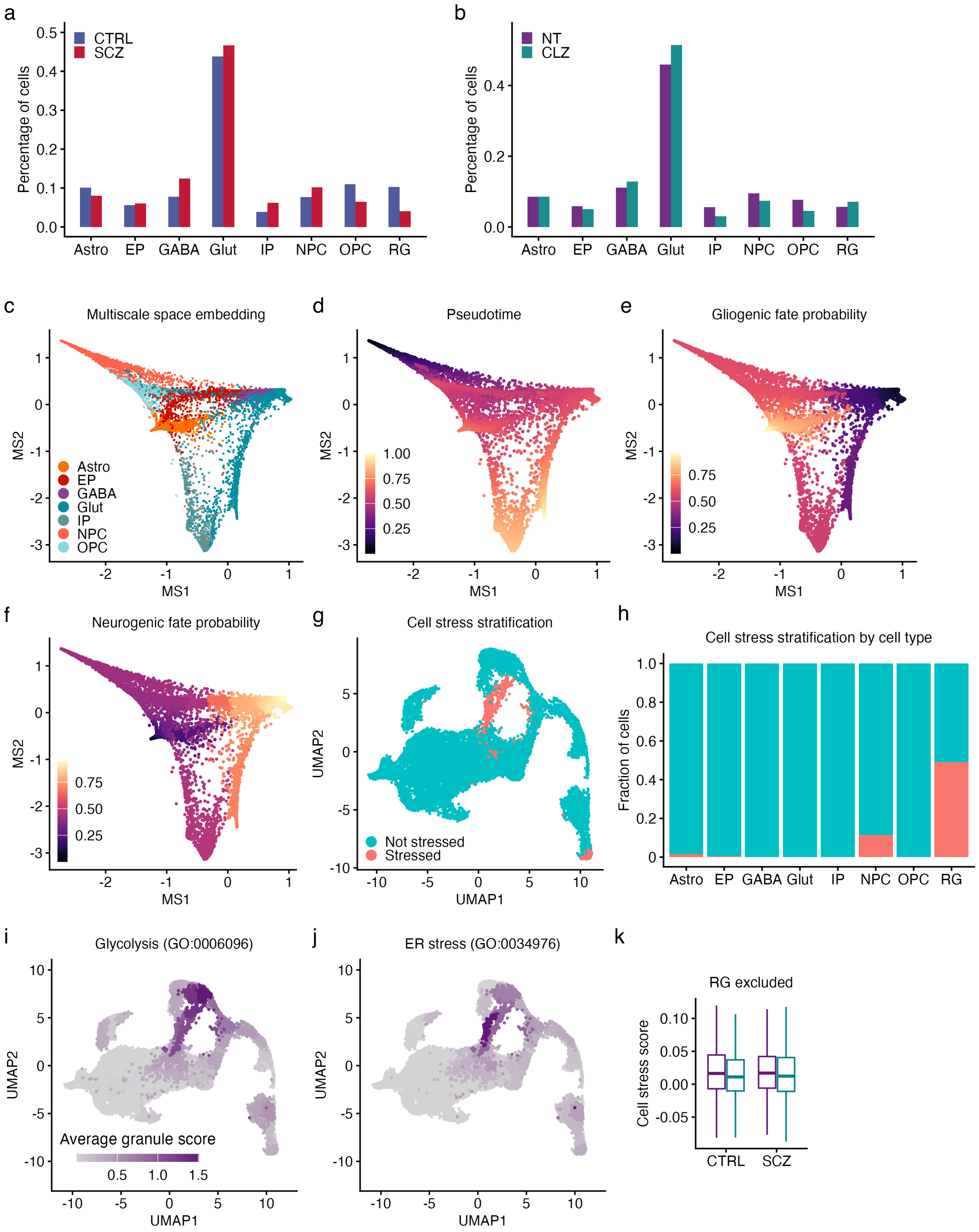


**Suppl. Figure 2. Pseudotime analysis and cell stress scoring of hCOs at single-cell resolution. a,b)** Cell type proportions by a) diagnosit and b) treatment conditions. No significant differences in donor-level cell type proportions were found using two-sided Wilcoxon tests. **b)** Multiscale space (MS) embeddings colored by hCO clusters. **d)** Pseudotime trajectories. Lighter color indicates increased differentiation from the initial NPC starting cell. **e,f)** Pseudotime analysis revealed two transcriptional trajectories corresponding e) gliogenic and f) neurogenic cell fates. **g)** Single cell annotation by cell stress status. Cell stress status was determined by automatic stress threshold estimation, defined as the theoretical 99% quantile of the fitted normal distribution. **h)** Cell stress stratification by cell types revealed approximately 50% overlap with the radial glia (RG) population. **i,j)** Average granule scoring based on expression of i) glycolysis and j) endoplasmic reticulum (ER) stress hallmark genes. **k)** RGs rely heavily on glycolysis for ATP production and may therefore confound cell stress quantification. Exclusion of RGs confirmed lack of CLZ-induced cellular stress. Paired two-sided Wilcoxon test on donor-level median stress scores.

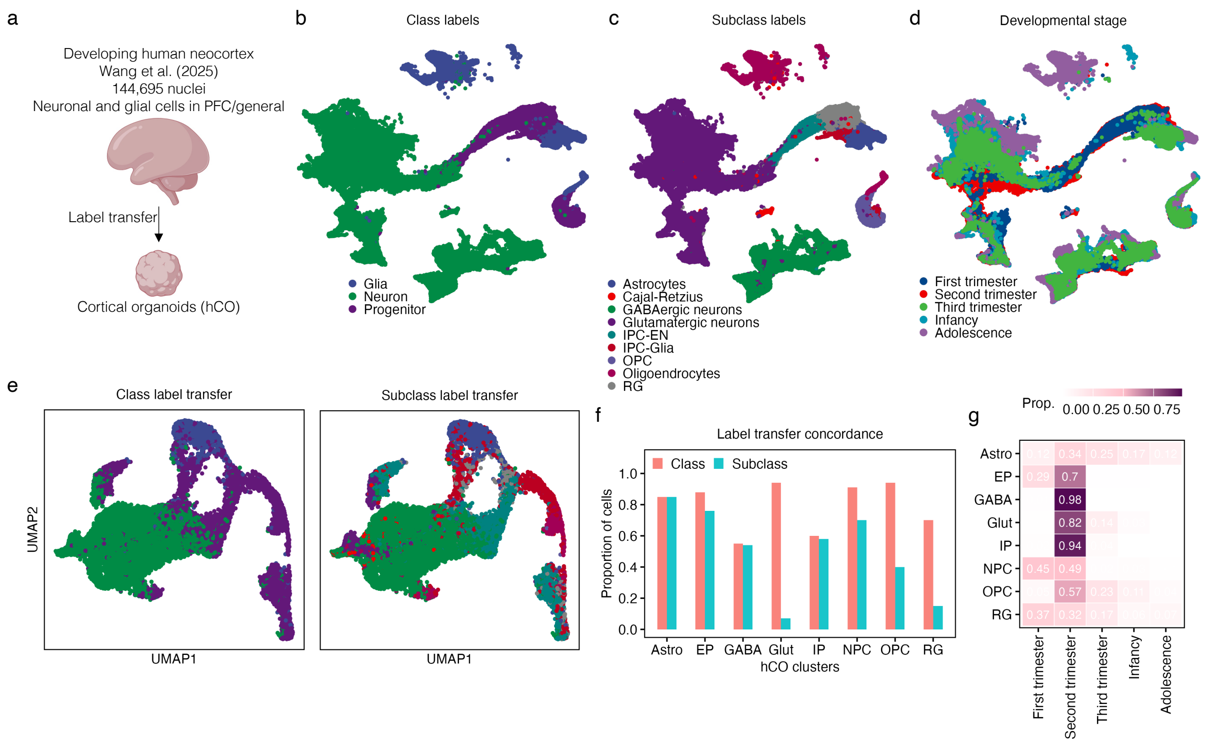


**Suppl. Figure 3. Cell label transfer from primary developing human neocortex to hCOs. a)** Neuronal and glial cells from prefrontal cortex (PFC) in the primary reference were considered for label transfer. **b)** UMAP embeddings of primary single-nucleus dataset colored by broad cell classes. **c)** UMAP embeddings colored by finer cell subclasses. IPC-EN, intermediate progenitors of excitatory neurons **d)** UMAP embeddings colored by developmental stages. **e)** Projection of primary cell classes and subclasses from primary dataset to hCOs. **f)** Concordance between assigned hCO cluster labels and transferred labels. **g)** Concordance between hCO profiles and developmental profiles from primary reference. Proportion indicates the percentage of cells in each hCO cluster showing highest transcriptional similarity with each developmental stage.

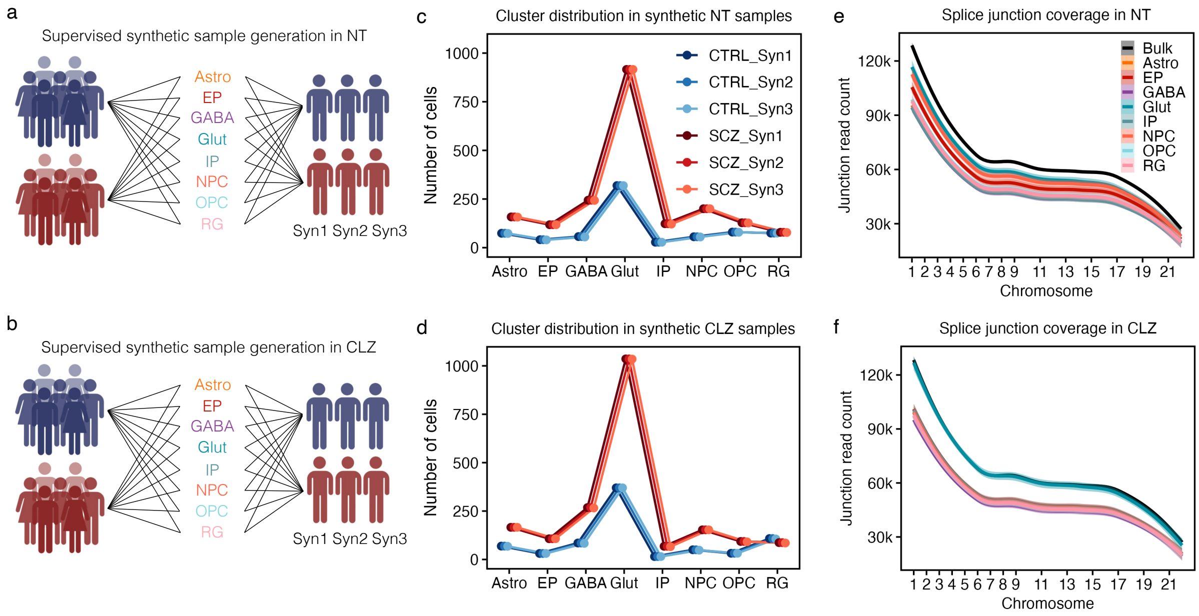


**Suppl. Figure 4. Synthetic sample generation in single-cell hCO data. a,b)** Overview of the synthetic sample generation strategy in a) non-treated (NT) and b) CLZ-treated hCOs. Within CTRL and SCZ samples, all cells of the same identity were first pooled together and then randomly distributed across three synthetic samples. This strategy resulted in 96 synthetic samples in total. **c,d)** Number of cells per cell type across synthetic samples. **e,f)** Cell type-specific splice junction coverage across synthetic samples. The coverage achieved was comparable to the coverage in the bulk RNA-seq data.

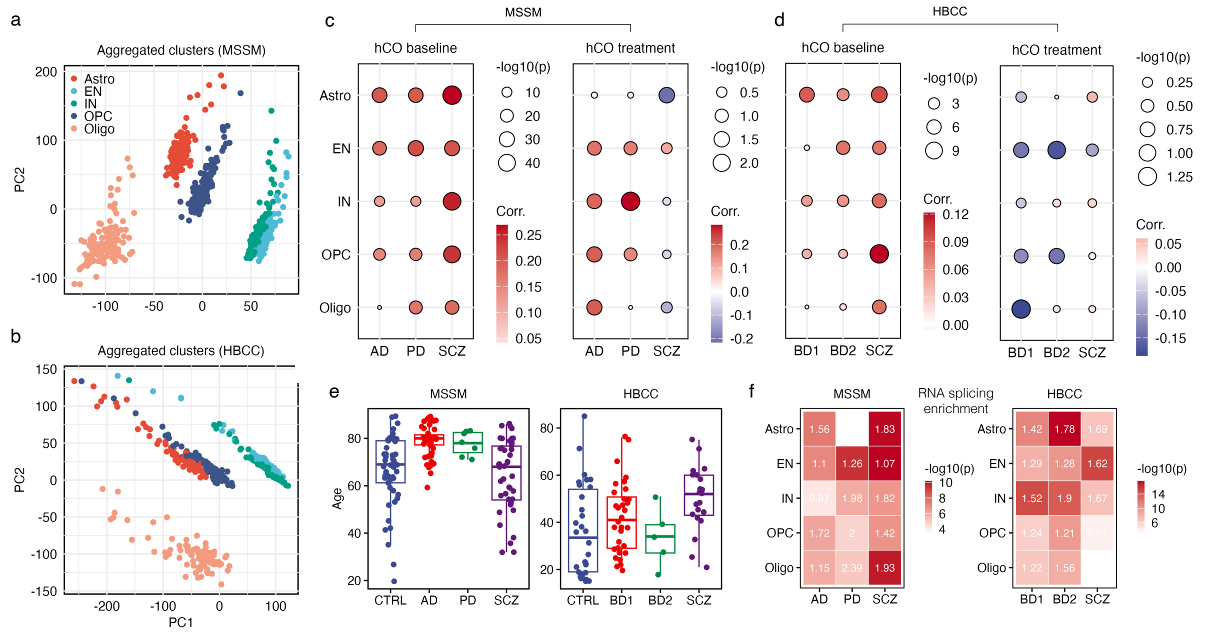


**Suppl. Figure 5. Projecting hCO signatures to primary disease profiles. a,b)** PCA of aggregated (pseudobulk) cell classes in a) MSSM and b) HBCC collections from the PsychAD primary disease reference. The plots show clear batch effects, particularly for the astrocyte and OPC clusters. **c,d)** Correlations between baseline and treatment hCO signatures and primary disease profiles in MSSM and HBCC collections across cell classes. **e)** Age distribution of postmortem donors in primary reference. In both collections, the age distribution of SCZ donors is comparable to the distribution in the other disorders, suggesting that the strong correlation between hCO signatures and primary SCZ profiles is little influenced by postmortem donor age. **f)** Enrichment of RNA splicing genes (annotated in GO) in primary disease profiles. SCZ is strongly enriched for RNA splicing genes in multiple cell types.

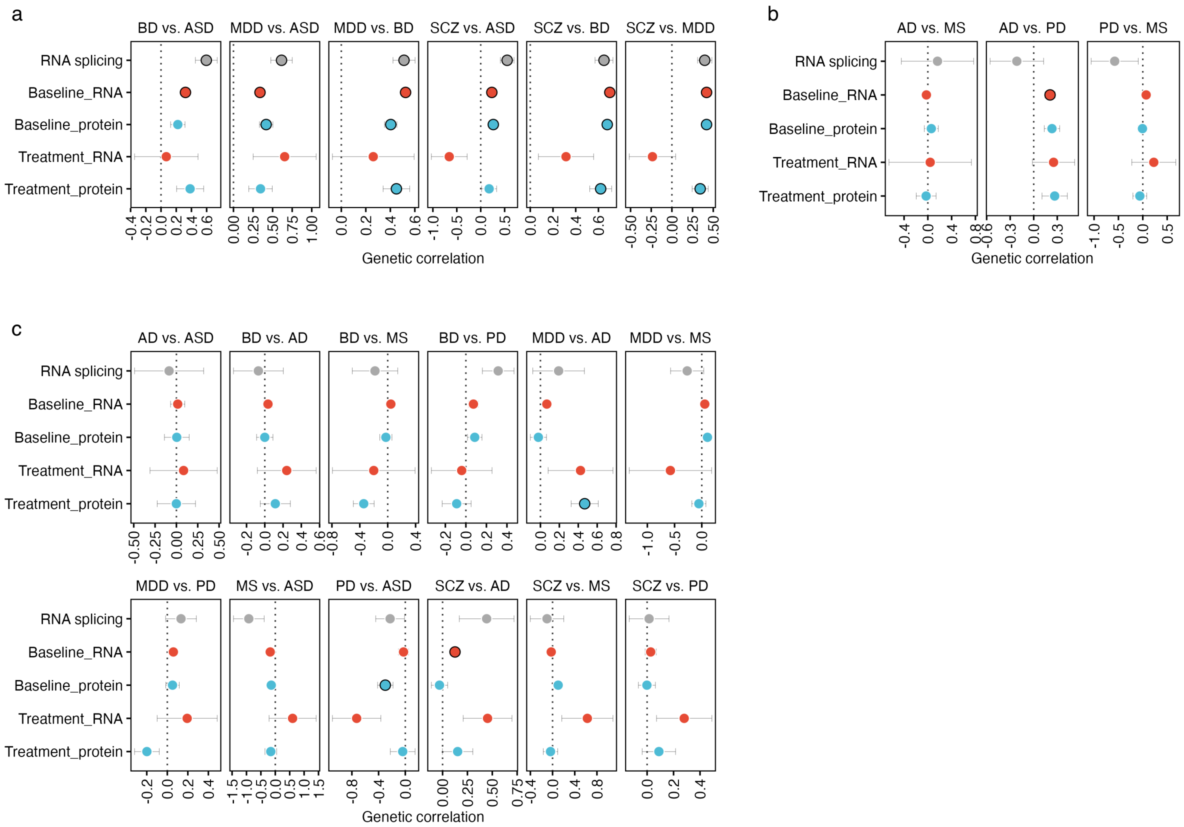


**Suppl. Figure 6. Bivariate genetic correlation of GWAS traits within RNA splicing and hCO signature gene sets. a)** Pairwise genetic correlations ($r_g$) between neuropsychiatric disorders. **b)** Genetic correlations among neurological disorders. **c)** Genetic correlations between neuropsychiatric and neurological disorders. Dots with black circles indicate significant (FDR < 0.05) genetic correlations. Most significant results were found between neuropsychiatric traits.

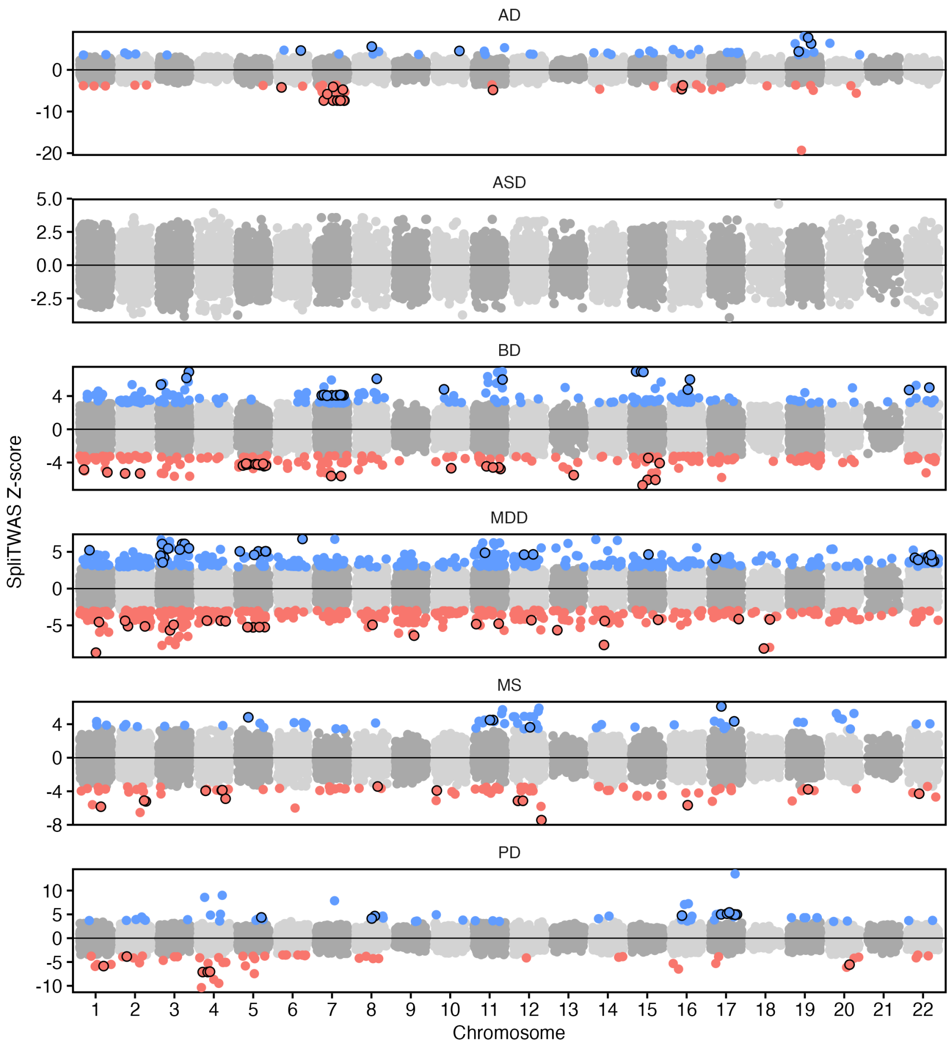


**Suppl. Figure 7. Splicing-specific TWAS (SpliTWAS) results.** The Manhattan-like plots show positive (Z-score > 0) and negative (Z-score < 0) associations in all disorders except for SCZ. Colored dots indicate significant associations (FDR < 0.05) and black circles indicate significant colocalization (posterior probability (PP4) > 0.8).
